# A quantitative demonstration of NADP^+^/NADPH redox homeostasis in cyanobacterial cells

**DOI:** 10.1101/2020.12.23.424211

**Authors:** Kenya Tanaka, Ginga Shimakawa, Hiro Tabata, Shoko Kusama, Chikahiro Miyake, Shuji Nakanishi

**Author notes:** Corresponding author: *Shuji Nakanishi*. **Email:**. **One sentence summary** Light-dependent photosynthetic reaction reduces only one third of total NADP^+^ in cyanobacterial cells, which limit the intracellular NADP^+^/NADPH ratio to a narrow range. **Author Contributions:** K.T., G.S., and S.N. designed research; K.T., G.S., and H.T. performed research; K.T., G.S., H.T., S.K. and S.N. analyzed data; and K.T., G.S., C.M., and S.N. wrote the paper.

## Abstract

In photosynthetic organisms, it is recognized that the intracellular NADP^+^/NADPH ratio is regulated within an appropriate range for the cooperative function of a wide variety of physiological processes. However, despite its importance, there is large variability in the values of the NADP^+^/NADPH ratio quantitatively estimated to date. In the present study, the light-response of the NADP^+^/NADPH ratio was investigated by applying a novel NADP(H) extraction method using phenol / chloroform / isoamyl alcohol (PCI) in the cyanobacterium *Synechocystis* sp. PCC 6803. The light-response of NADP(H) observed using PCI extraction was qualitatively consistent with the NADPH fluorescence time course measured *in vivo*. Moreover, the results obtained by PCI extraction and the fluorescence-based methods were also consistent in a mutant lacking the ability to oxidize NAD(P)H in the respiratory chain, and exhibiting a unique NADPH light-response. These observations indicate that the PCI extraction method allowed quantitative determination of NADP(H) redox. Notably, the PCI extraction method showed that not all NADP(H) was oxidized or reduced by light-dark transition, indicating that some NADP(H) is not light-responsive. Specifically, 64% of total NADP(H) was observed as non-light-responsive in the wild-type cells. The variation of the intracellular NADP^+^/NADPH ratio is limited to a narrow range due to the presence of non-light-responsive NADP(H).

## Introduction

The redox pair NADP^+^/NADPH is involved in various reactions in photosynthetic organisms. NADP^+^ is the terminal electron acceptor in the photosynthetic electron transport chain (PETC) and is converted to NADPH under light conditions. The NADPH thus generated supports biosynthetic and antioxidant systems by serving as a reducing driver for various enzymes including glyceraldehyde 3-phosphate dehydrogenase (GAPDH), NADPH-thioredoxin reductase (NTR), glutathione reductase (GR), etc (Raines, 2003; Hishiya et al., 2008; Yoshida and Hisabori, 2016; Vogelsang and Dietz, 2020). The oxidized form (NADP^+^) also functions as an essential co-factor for glucose-6-phosphate dehydrogenase (G6PDH), 6-phosphogluconate dehydrogenase (6PGDH), and isocitrate dehydrogenase (ICD) in primary metabolism (Muro-Pastor and Florencio, 1992; Ishikawa and Kawai-Yamada, 2019). Given that the NADP^+^/NADPH redox ratio, a critical factor influencing various biological processes, can vary with changes in the lighting conditions, it is vitally important to quantitatively determine the intracellular concentrations of NADP(H) for a deeper understanding of the physiology of photosynthesis.

Fluorescence detection of NADPH is a representative *in vivo* method for measuring light-responsive changes in NADPH concentrations. For example, it has been shown that NADPH was produced or consumed in the sub-second order by light-dark transitions (Mi et al., 2000; Kauny and Sétif, 2014; Shaku et al., 2016). However, the fluorescent yield of NADPH changes depends on the peripheral environment (Latouche et al., 2000; Kauny and Sétif, 2014). Importantly, NADP^+^ (the counterpart of NADPH) does not emit fluorescence. Therefore, the *in vivo* fluorescence-based technique cannot directly assess the NADP^+^/NADPH redox ratio or the absolute amount of NADP(H). In contrast, the concentration of NADP(H) can be quantitatively measured following cellular extraction. In fact, previous studies have attempted to quantify the concentration of NADP(H) and the redox ratio *in vitro*. However, the reported values of the *in vitro* studies differ widely (Table 1). The redox ratio estimated by *in vitro* extraction methods must be consistent with the dynamic behavior of NADPH fluorescence; however, no studies have been undertaken to demonstrate consistency between *in vivo* and *in vitro* measures.

**Table 1.**
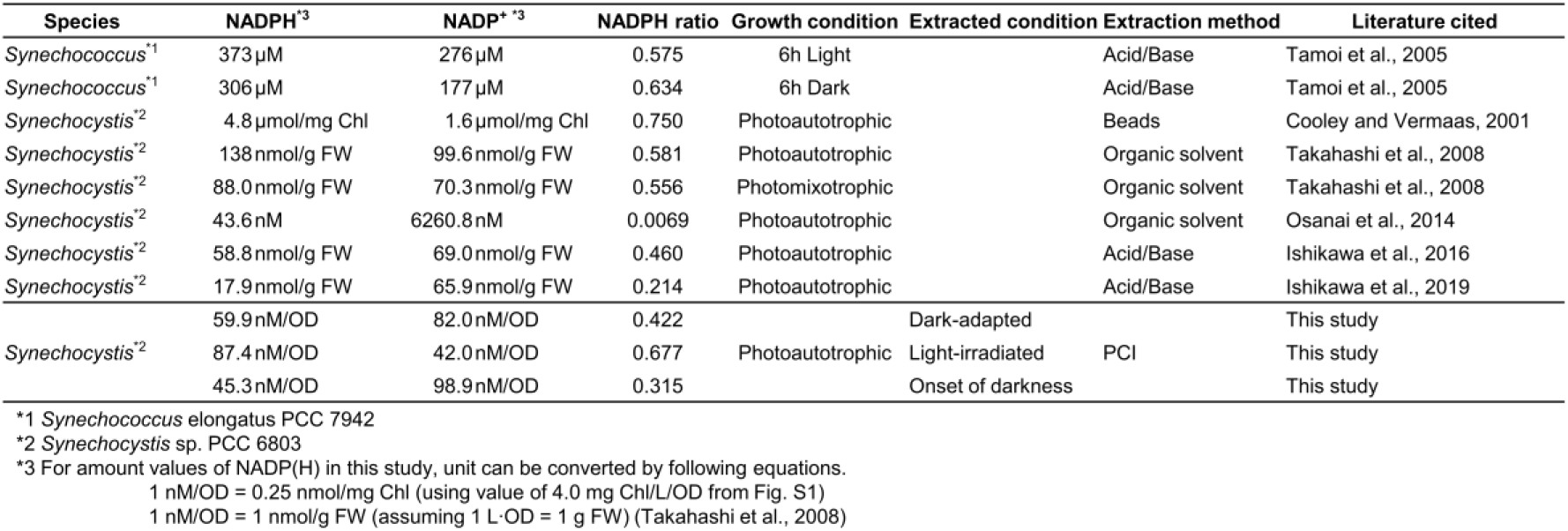
Comparison of NADP(H) measurements in cyanobacteria and their results.

In the present work, we quantified the amount of NADP(H) in *Synechocystis* sp. PCC 6803 (hereafter *Synechocystis*) *in vitro* using a phenol/chloroform/isoamyl alcohol (PCI) solution to deactivate undesirable enzymatic reactions, and evaluated the obtained values against the dynamic behavior of NADPH fluorescence. Importantly, the light-responsive changes in NADP(H) determined using the PCI-extraction method were consistent with the results of *in vivo* NADPH fluorescence measurements, indicating the validity of the novel extraction protocol. Moreover, development of this novel protocol has revealed the novel finding that 64% of the NADP^+^/NADPH redox pair in *Synechocystis* was unchanged by light-dark transition, i.e., non-light-responsive. Due to the presence of the non-light-responsive NADP(H) pool, the redox potential of intracellular NADP^+^/NADPH varies only 14 mV between light and dark conditions.

## Results

### *In vitro* quantitation of NADP(H)

First, NADP(H) was extracted from dark-adapted *Synechocystis* cells using extraction buffer, and subsequently quantified according to the standard protocol of the commercially-available assay kit (see Materials and Methods for details). Although 2.1 μM of NADP+ in the extraction buffer was confirmed to be present, the reduced form, NADPH, was not detected (Fig. 1A-1). To verify the possibility that NADPH was unexpectedly oxidized during the extraction process, extraction was performed using a buffer containing exogenous 1.5 μM NADP+ and 1.5 μM NADPH as standards. The NADPH level was confirmed to be less than 1.5 μM even for this sample (Fig. 1A-2). Notably, when only the exogenous NADP+/NADPH mixture was examined, the concentrations were accurately quantified (Fig. 1A-3). These results clearly indicated that components in the cell lysate oxidized NADPH during the extraction process.

**Figure 1.**
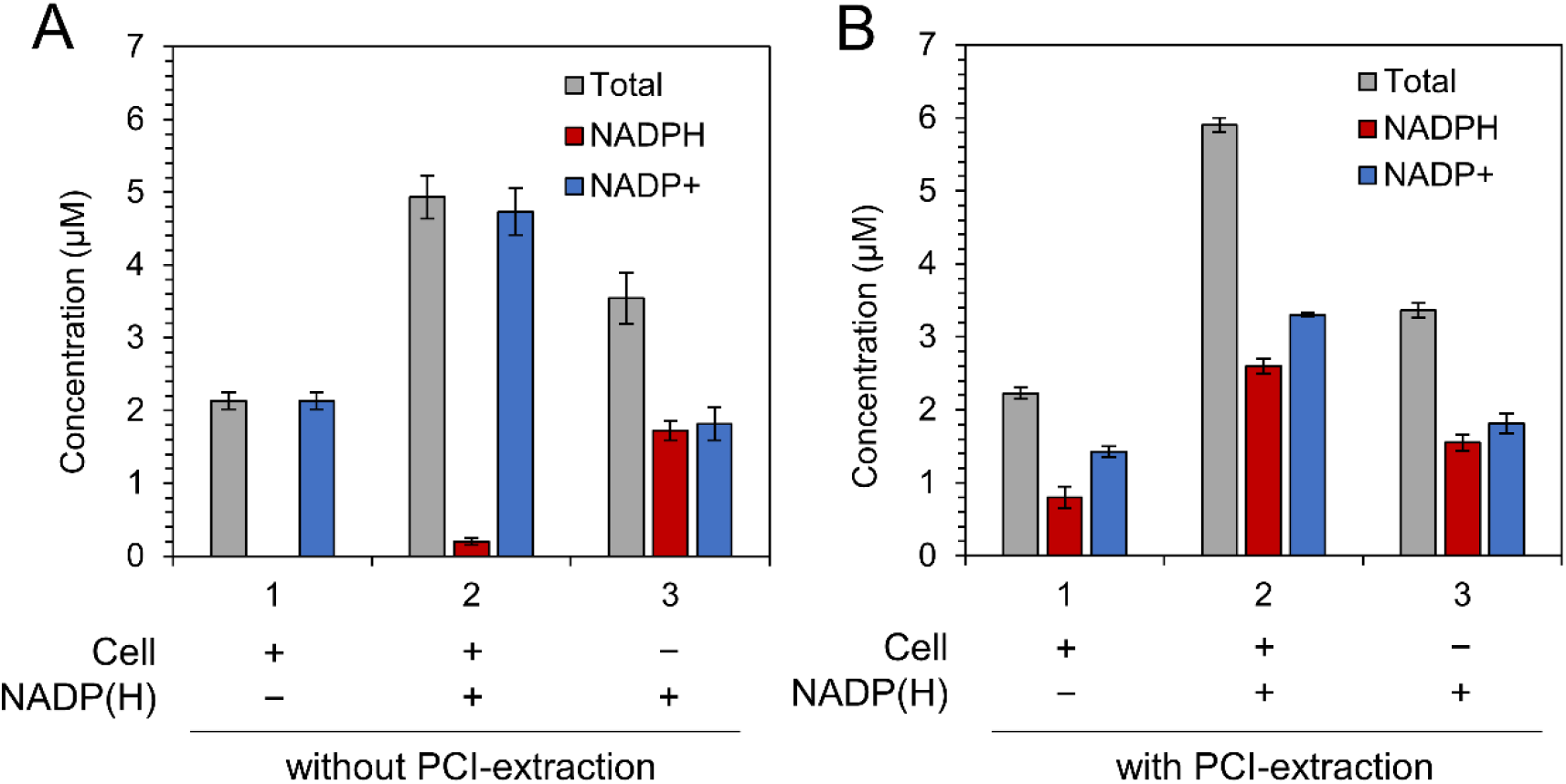
Effect of PCI solution on NADP(H) extraction. NADP(H) concentration in the crude extracts processed (A) without and (B) with PCI. Sample numbers are indicated on the horizontal axes; No. 1: crude extracts from dark-adapted cells, No. 2: crude extracts containing exogenous NADP(H), No. 3: exogenous NADP(H). Values are means ± SD (bars) of three biological replicates.

Considering that non-enzymatic degradation of NADP(H) is negligible due the very slow kinetics, Fig. 1A suggests that the cell lysate contains enzymes that accelerate the redox reactions related to NADP(H) chemistry. To accurately quantify the intracellular NADP^+^/NADPH ratio, these proteins must be quickly inactivated during the extraction process. Therefore, NADP(H) was extracted in the presence of a PCI, a protein scavenging agent widely used for extracting nucleic acids (hereafter, this protocol is called PCI-extraction). In this case, although the total concentration of NADP^+^ and NADPH was the same as that determined without PCI-treatment, the NADPH value reached 0.8 μM (Fig. 1B-1). When 1.5 μM of exogenous NADP(H) was added following addition of PCI to the cells, the concentrations of NADPH and NADP^+^ both increased by approximately 1.5 μM (Fig. 1B-2). It was confirmed that PCI itself did not affect the measurement of exogenous NADP(H) (Fig. 1B-3). These results suggested that the intracellular concentrations of NADP(H) can be quantitatively determined using the developed PCI-extraction protocol.

### Comparison of *in vivo* and *in vitro* measurements

To evaluate the validity of the PCI-extraction protocol, the time-transient behavior of NADPH levels under varying irradiation conditions was compared with the results obtained using the well-established NADPH fluorescence measurement method. As shown in Figure 2A, when the actinic light was turned on, the NADPH fluorescence intensity rapidly increased (stage-II). When the light was turned off, the fluorescence intensity decreased and reached a minimum within 5 s, subsequently increasing and stabilizing after 30 s (stage-III). This time-transient behavior is in good agreement with previous reports, and the decrease and increase of the fluorescence level after turning the light off in stage-III are considered to be due to NADPH consumption by the Calvin cycle and NADPH production by the oxidative pentose phosphate pathway (OPPP), respectively (Mi et al., 2000; Kauny and Sétif, 2014). The fluorescence-based method can detect NADPH responses to environmental light changes on a time scale of seconds. In this study, we define the NADP(H) that can be detected by fluorescence, corresponding to the orange zone in Fig. 2A, as “light-responsive NADP(H)”.

**Figure 2.**
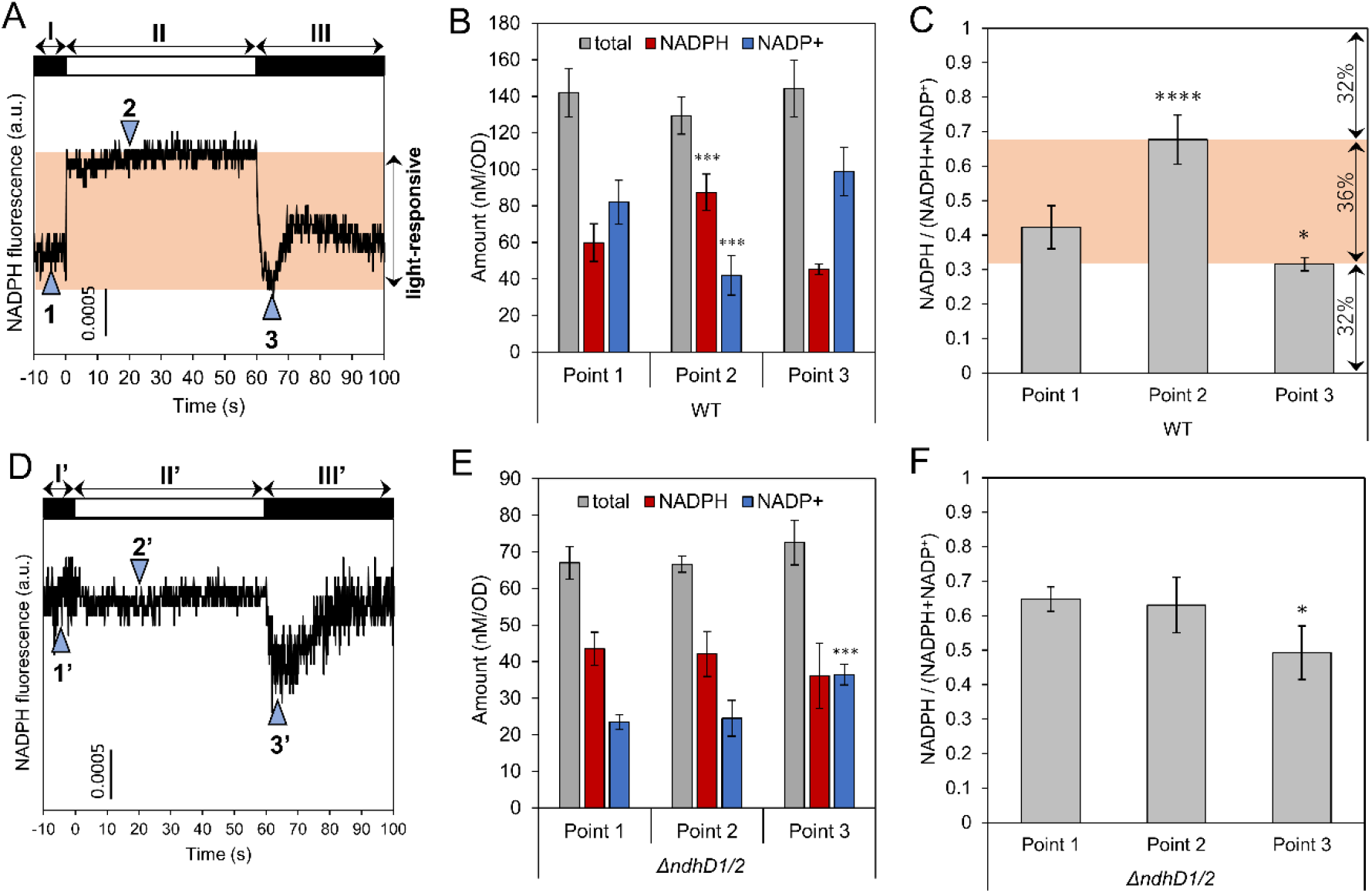
Light-response of NADP(H) in (A-C) WT and (D-F) *∆ndhD1/2* mutant. (A, D) NADPH fluorescence transients induced by 1 min illumination. White and black bars indicate light and dark condition, respectively. Light-responsive NADP(H) in WT is indicated by the orange zone. Points indicated by triangles with numbers correspond to three different states; point 1: after 1 h dark adaptation, point 2: 20 s after light irradiation, point 3: 5 s (3 s for *∆ndhD1/2*) after transition to the dark condition. (B, C, E, F) NADP(H) contents and NADPH ratio against total NADP(H) content in WT (B, C) and *∆ndhD1/2* (E, F) obtained by adding PCI to the cell suspension at each time point. Range of light-responsive NADP(H) ratio in WT is indicated by the orange zone. Values are means ± SD (bars) of about 3 ‒ 6 biological replicates. Significant differences from dark-adapted conditions were evaluated by a student’s t test (*P < 0.05, *** P < 0.001, **** P < 0.0001).

Next, the range of variation in the amount of NADP(H) was estimated using the PCI-extraction method. PCI was added to cell suspension at three different points (Fig. 2A, points 1-3). Although the total amounts of NADP(H) were the same for all three samples, the NADP^+^/NADPH ratio changed depending on the stage (Figs. 2B, C). The observed relative ratio of NADPH was 0.42 for the dark-adapted cells, which increased to 0.68 following light irradiation, and then decreased to 0.32 at the onset of darkness. The time-transient behavior of NADPH levels under light/dark perturbations deduced using the PCI extraction protocol agrees well with that obtained using fluorescence measurements. It is noteworthy that the ratio of NADPH to total NADP(H) at points 2 and 3, in which the fluorescence gave the maximum and minimum values, was not 100% and 0%, respectively. This is an important observation, since it indicates that some of the NADP(H) is not light-responsive. In this experimental condition, the relative amounts of light-responsive NADPH, non-light-responsive NADP^+^, and non-light-responsive NADPH were estimated to be 36%, 32% and 32%, respectively (Fig. 2C).

The above conclusion was further verified using a mutant strain lacking *ndhD1* and *ndhD2* (*∆ndhD1/2*). In this mutant strain, which lacks the ability to oxidize NAD(P)H in the respiratory chain through the Type I NAD(P)H dehydrogenase complex (NDH-1) (Ohkawa et al., 2000), the relative ratio of NADPH at the dark-adapted state is known to be as high as that in the light-adapted state (Sétif et al., 2020). In fact, NADPH fluorescence levels did not change when the light was turned on (Fig. 2D, stages I’ and II’). Although the fluorescence level decreased transiently after the light was turned off, it quickly recovered to the same level as in the light condition (stage III’). This behavior was consistent with previous results (Sétif et al., 2020). The PCI-extraction method was then applied to the mutant strain for comparison with the results of NADPH fluorescence measurements. For the mutant strain, there was no significant difference in the relative ratio of NADPH between the dark adapted and light irradiated points (Figs. 2E and F, points 1’ and 2’), whereas a decrease in the NADPH ratio was observed 3 s after the light was turned off (point 3’). This behavior is consistent with that obtained by the NADPH fluorescence.

Kauny et al. reported that the amount of light-dependent NADPH (difference between points 1 and 2 in Fig. 2A) was overestimated in the fluorescence method, based on calibration curves obtained using exogenous NADPH (Kauny and Sétif, 2014). More specifically, due to the higher fluorescence of protein-bound intracellular NADPH, the enhancement factor reaches 2 to 4. In our results, light-dependent NADPH was estimated to be 20.9 and 9.0 nmol mg^−1^ Chl using the fluorescence-based and PCI-extraction methods, respectively (Figs. 2A and S2). Thus, the enhancement factor was 2.3, which is within the range reported by Kauny et al. Notably, assuming that all NADPH (35.5 nmol mg^−1^ Chl) is light-responsive, the enhancement factor is calculated to be less than 1, which is inconsistent with the previous report. These results also supported the existence of non-light-responsive NADP(H).

## Discussion

As described, our experimental results clearly show that some of the NADP(H) pool is not light-responsive. As shown in Fig. 2C, light-responsive NADP(H), non-light-responsive NADP^+^ and non-light-responsive NADPH accounted for about one-third of the total amount of NADP(H) in wild-type (WT) cells. Since NADP^+^ is an essential cofactor for G6PDH, 6PGDH and ICD, which are components of primary metabolism pathways, some portion of NADP^+^ pool needs to be stably present even under light conditions, i.e., non-light-responsive. On the other hand, NADPH provides essential reducing power for maintaining antioxidant ability through enzymes such as NTR and GR under both light and dark conditions (Hishiya et al., 2008; Yoshida and Hisabori, 2016; Vogelsang and Dietz, 2020). Thus, given that NADP^+^ and NADPH have their own distinctive roles in photosynthetic cells, maintaining an appropriate balance between the non-light-responsive and light-responsive NADP(H) pool is expected to be important in facilitating various biological processes cooperatively in fluctuating light environments. For plastoquinone (PQ), one of the major intracellular redox species, it is reported that photoactive and non-photoactive pools exist in both chloroplasts and cyanobacteria (Kruk and Karpinski, 2006; Khorobrykh et al., 2020). In fact, we found that non-photoactive PQ accounted for 87% of the PQ pool in *Synechococcus elongatus* PCC 7942 cells (Fig. S3). Although NADP(H) is located in the cytoplasm, and cyanobacteria lack organelles, light-responsive and non-light-responsive NADP(H) can be partitioned in a cyanobacterial cells by an unknown mechanism. For example, it was reported that NADPH allosterically decreases the binding affinity of ferredoxin (Fd) to ferredoxin-NADP+ reductase (FNR) (Kimata-Ariga et al., 2019), which may prevent complete reduction of the NADP^+^ pool. Another possible explanation is that NADP(H) nonspecifically binds proteins such as Rubisco (Badger and Lorimer, 1981; Latouche et al., 2000), which may represent the bulk of the non-light-responsive NADP(H) pool. Furthermore, the possibility that some proportion of NADP(H) is spatially sequestered into unknown compartments in cyanobacterial cells cannot be ruled out. Thus, there is room for further investigation toward unveiling the nature of the non-light-responsive NADP(H) pool.

Next, let us consider the reasons for the increase and decrease of the light-responsive NADP(H) pool. It is obvious that (light-responsive) NADPH is increased by the reduction of NADP^+^ at the PETC under illumination (Mi et al., 2000; Kauny and Sétif, 2014; Shaku et al., 2016). The initial decay in NADPH fluorescence just after the onset of the dark phase (Fig. 2A, before point 3 in stage III) can be explained by the increased consumption of NADPH in the Calvin cycle (Kauny and Sétif, 2014). The subsequent increase in fluorescence intensity (after point 3) is thought to be due to NADPH production by increased OPPP activity. On the other hand, the *∆ndhD1/2* strain exhibited different time-transient behavior in fluorescence intensity from that observed for the WT (Fig. 2D), indicating that not only the PETC but the respiratory electron transport chain can also lead to light-responsive change in NADP(H) amount. Moreover, as shown in Fig. 2E, the total amount of NADP^+^ and NADPH for the mutant was approximately half of that for the WT. Taken together, activities in the photosynthetic and respiratory electron transport chains, Calvin cycle, and OPPP contribute to the amount of light-responsive NADP(H).

Figure 3A shows the Nernstian relationship between the NADP^+^/NADPH redox ratio and the redox potential, which was drawn based on the reported value of the standard redox potential, −320 mV (Rodkey and Donovan, 1959; Burton, 1974; Foyer and Noctor, 2016). The mid-point potential (*E*_M_) for enzymes related to NADP(H) chemistry are also shown in the same figure as reference information (Keirns and Wang, 1972; Foyer and Noctor, 2016; Mihara et al., 2020). Based on the NADP^+^/NADPH ratio quantitatively estimated in this work (Fig. 2), the redox potential of NADP^+^/NADPH was estimated to be −316 (± 3) mV for the dark-adapted state cells and −330 (± 4) mV for the irradiated cells. These potentials must be higher than the *E*_M_ of enzymes with NADP^+^ as the electron acceptor and lower than the *E*_M_ of enzymes with NADPH as the electron donor. In fact, the redox potentials were respectively higher and lower than the *E*_M_ of FNR catalyzing the reduction of NADP^+^ (−360 mV) (Keirns and Wang, 1972) and thioredoxin needed for reductive quenching of reactive oxygen species (approximately −304 to −257 mV) (Mihara et al., 2020). Figure 3B shows the redox states of various redox enzymes based on the Nernstian relationship, under the assumption that intracellular redox is globally governed by the redox potential of NADP^+^/NADPH. Although we cannot make a rigorous discussion based on equilibrium, the equilibrium potential of the NADP^+^/NADPH pair can be a useful indicator of the global redox state in cells, since NADP(H) is one of the major intracellular redox species (Foyer and Noctor, 2016) and NADPH generated at the PETC diffuses rapidly throughout the cell (see also supporting information). As can be seen in Fig. 3B, FNR was maintained in an oxidative state, whereas antioxidant enzymes such as thioredoxin were in the reductive state, under both light and dark conditions. Thus, it was shown from our quantitative analyses that both the suppression of electron accumulation in the PETC and the stable supply of electrons to the antioxidant systems are indeed thermodynamically allowed.

**Figure 3.**
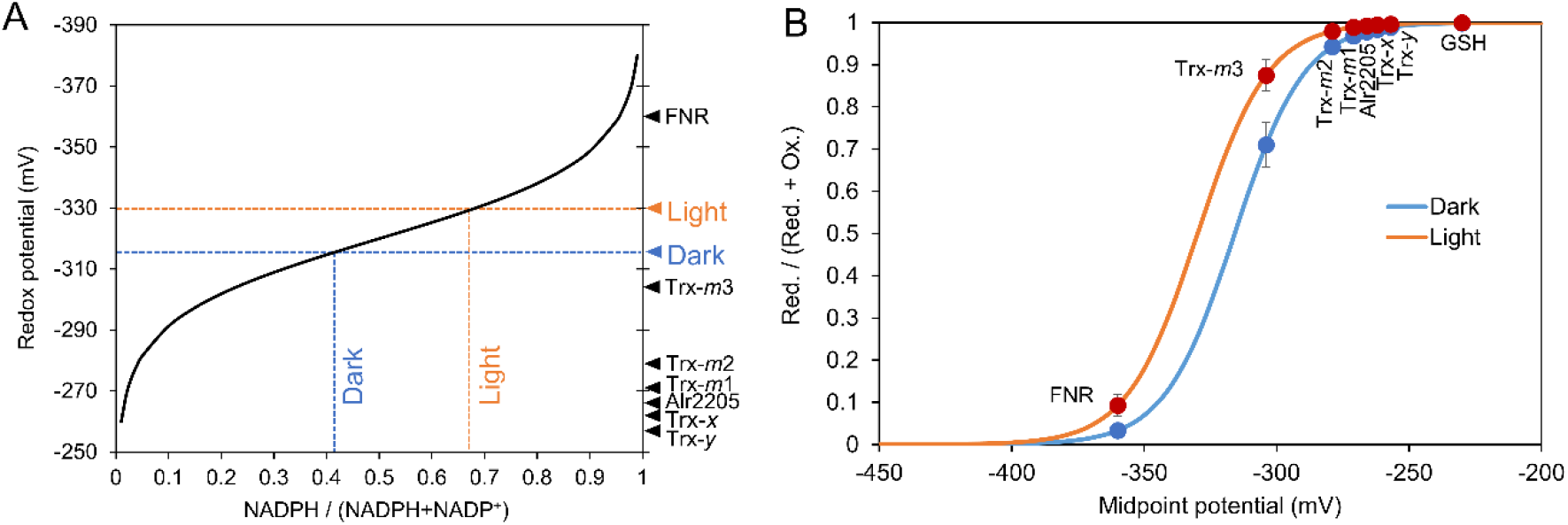
Difference in redox potential between dark and light conditions. (A) Redox potential of NADP(H) in dark and light conditions are respectively calculated using the experimental values shown in Fig. 2C, and indicated as blue and orange lines with the theoretical Nernst curve (black solid line). Mid-point potentials of enzymes related to NADP(H) are shown on the right vertical axis. (B) Ratio of reduced (Red.) and oxidized (Ox.) forms of redox species under a condition where the redox potential of NADP(H) prevails so that all the components are the same potential value is shown based on the Nernst relation. Blue and orange lines indicate Nernst curves under the redox potential of −316 mV (dark condition) and −330 mV (light condition). Values are means ± SD (bars) of 6 results calculated from the biological replicates in Fig. 2C.

As described, we have successfully determined the absolute amount of NADP(H) and the NADP^+^/NADPH ratio using PCI during the extraction process. The results obtained using this *in vitro* method were in very good agreement with the results obtained from the *in vivo* fluorescence method, demonstrating that this treatment is capable of halting cellular redox within at least a few seconds. This capability is likely to be attributed to the quick inactivation of proteins by phenol, which prohibits further changes in the redox state of NADP(H). Importantly, our quantitative analyses revealed that the redox potential of NADP^+^/NADPH varies by only 14 mV between light and dark conditions, due to the presence of non-light-responsive NADP(H). This is the first quantitative demonstration of the redox homeostasis of NADP(H). We anticipate further quantitative studies aiming to clarify the separation mechanisms and roles of light-responsive and non-light-responsive NADP(H) will deepen our understanding of the regulation mechanisms and physiology of photosynthesis.

## Materials and Methods

### Bacterial strains and cell culture conditions

We used the following *Synechocystis* strains: wild-type and *∆ndhD1/2* (Ohkawa et al., 2000). These strains were grown and maintained on solid (1.5% Bacto agar) BG-11 medium plates. For pre-culture, cells from the agar plate were inoculated into 30 mL liquid BG-11 medium in a 100-mL flask and grown at 30 °C with air bubbling under white light illumination at an intensity of 20 μmol m^−2^ s^−1^. For the main culture, the pre-culture was inoculated to achieve an optical density of 0.02 at 730 nm (OD_730_) in 30 mL BG11. Other conditions were the same as those in pre-culture. Fig. S1 shows the growth curves of the main culture measured at an optical density of 730 nm and the chlorophyll concentration. To determine the chlorophyll concentration, the cells were harvested by centrifugation at 12,000 × g for 5 min and suspended in 100% methanol. The suspension was then centrifuged at 12,000 × g for 5 min, followed by measurement of absorption at 665 nm. The chlorophyll *a* concentration was calculated according to a previously described method (Grimme and Boardman, 1972).

### NADPH fluorescence measurements

The *in vivo* NADPH fluorescence originating from NAD(P)H was measured using the NADPH/9-AA module of a Dual-PAM-100 instrument (Heinz Walz, Effeltrich, Germany) (Kauny and Sétif, 2014; Shimakawa et al., 2018). The reaction mixtures (2 mL) contained fresh BG-11 medium (pH 7.5) and cyanobacterial cells (2.5 μg chlorophyll mL^−1^). The NADPH/9-AA module consists of an emitter unit (DUAL-ENADPH) and a detector unit (DUAL-DNADPH). NADPH fluorescence was excited by UV-A (365 nm) from the DUAL-ENADPH unit and detected by a blue-sensitive photomultiplier with a filter transmitting light between 420 and 580 nm in the DUAL-DNADPH unit. The measured light intensity was on a scale from 1 to 20, and the intensity was set at 10 in this study. The measured light frequency in the absence and presence of red actinic light (200 μmol photons m^−2^ s^−1^) was set at 200 Hz and 5,000 Hz, respectively.

### NADP(H) extraction

PCI solution was prepared as follows. Crystalline phenol was melted in a 65 °C water bath, followed by the addition of an equal volume of 0.5 M Tris-HCl buffer (pH 8.0). After vigorous mixing, the upper water phase was removed and 0.1 M Tris-HCl (pH 8.0) was added. The same amount of chloroform/isoamyl alcohol (24:1 v/v) solution as the phenol was added to obtain the PCI solution (phenol : chloroform : isoamyl alcohol 25:24:1 v/v). NADP(H) extraction was performed using about 5 to 6 day-old main cultures corresponding to OD_730_ = 2 ‒ 4 (approx.) (Fig. S1). Before extraction, the cultures were maintained under darkness for an hour with air bubbling. After the dark adaptation, cells were harvested from the calculated volume of culture (2 ml / OD_730_) by centrifugation at 12,000 × g for 3 min. The cell pellet was suspended in 50 μL of 1 mM NaHCO_3_. For the extraction from dark-adapted cells, 300 μL PCI was added to the suspension followed by the addition of 250 μL extraction buffer (approx. pH 10.4 ‒ 11.0, production code: N510 Dojindo, Kumamoto Japan). For light-irradiated extraction, the cell suspension was transferred into a cuvette with 1 mm optical path to illuminate the cells. LED light of 600 μmol m^−2^ s^−1^ (pE-100wht, BioVision Technologies, Exton, PA, USA) was used for actinic light. A 300-μL aliquot of PCI was added to the suspension in the cuvette at various time points, followed by the addition of 250 μL extraction buffer. The PCI suspension samples were centrifuged at 12,000 × g for 3 min. Each upper water phase was transferred to a microtube and frozen quickly in liquid nitrogen.

For extraction without PCI, only extraction buffer was added to the cells. The cell suspension was frozen and thawed once for cell-disruption. Proteins in the cell lysate were removed by ultrafiltration before NADP(H) measurements. Other extraction processes were same as those for PCI.

### Enzymatic NADP(H) measurement

For the PCI extraction samples, after thawing the NADP(H) extraction, 250 μL chloroform/isoamyl alcohol (24:1 v/v) solution was added for further removal of phenol. The mixture was centrifuged at 12,000 × g for 5 min and the water layer was transferred into two microtubes for measurement of total NADP(H) and NADPH. The sample used for measuring NADPH was incubated at 60 °C for an hour to decompose NADP^+^ in the sample solution (NADP^+^ is unstable in heated alkaline solutions), while the sample used for measuring total NADP(H) was kept on ice. For determination of the NADP(H) concentration, the enzymatic cycling assay was performed according to the manufacturer’s instructions (production code: N510 Dojindo). Absorbance at 450 nm of each sample in a 96-well plate was measured on Infinite M200 Plate Reader (Tecan, Männedorf, Switzerland).

## Acknowledgments

We thank Dr. Ohkawa (Hirosaki University) and Prof. K. Sonoike (Waseda University) for kindly providing the mutant used in this work. This work was partially supported by the Advanced Low Carbon Technology Research and Development Program (JPMJAL1402) of the Japan Science and Technology Agency (JST), and JSPS KAKENHI Grant Number 18J20176 and 20J00105.

